# Predicting Evolutionarily Stable Strategies from Functional Responses of Sonoran Desert Annuals to Precipitation

**DOI:** 10.1101/271080

**Authors:** William S. Cuello, Jennifer R. Gremer, Pete C. Trimmer, Andrew Sih, Sebastian J. Schreiber

**Affiliations:** University of California, Dept. of Mathematics, 1 Shields Avenue, Davis, CA 95616; University of California, Dept. of Evolution and Ecology, 1 Shields Avenue, Davis, CA 95616; University of California, Dept. of Environmental Science and Policy, 1 Shields Avenue, Davis, CA 95616

**Keywords:** Bet-hedging, Desert Annuals, Germination, Water-Use Efficiency, Evolutionarily Stable Strate-gies, Seed Survival

## Abstract

For many decades, researchers have studied how plants use bet-hedging strategies to insure against unpredictable, unfavorable conditions. We improve upon earlier analyses by explicitly accounting for how variable precipitation affects annual plant species’ bet-hedging strategies. We consider how the survival rates of dormant seeds (in a ‘seed bank’) interact with precipitation responses to influence optimal germination strategies. Specifically, we incorporate how response to resource availability (i.e. the amount of offspring (seeds) generated per plant in response to variation in desert rainfall) influences the evolution of germination fractions. Using data from 10 Sonoran Desert annual plants, we develop models that explicitly include these responses to model fitness as a function of precipitation. For each of the species, we identify the predicted evolutionarily stable strategies (ESS) for the fraction of seeds germinating each year and then compare our estimated ESS values to the observed germination fractions. We also explore the relative importance of seed survival and precipitation responses in shaping germination strategies by regressing ESS values and observed germination fractions against these traits. We find that germination fractions are lower for species with higher seed survival, with lower reproductive success in dry years, and with better yield responses in wet years. These results illuminate the evolution of bet-hedging strategies in an iconic system, and provide a framework for predicting how current and future environmental conditions may reshape those strategies.

## Introduction

For organisms inhabiting variable environments, it can be difficult to perfectly time key life history functions, such as emergence or reproduction, with favorable conditions. In such environments, natural selection will favor traits or strategies that buffer against uncertainty, usually by spreading risk over time or space. This is often achieved through bet-hedging, in which strategies or traits reduce variance in reproductive success at some cost to mean reproductive success [1–3]. Empirical examples of putative bet hedging strategies include iteroparity [4], variable diapause [5, 6], and variable offspring size [7, 8]. Delayed seed germination in annual plants is the classic example of bet-hedging [9], in which variance in success is reduced by spreading germination over multiple years, which has been clearly demonstrated in annual plants of the Sonoran Desert [10, 11].

Bet-hedging strategies interact with other traits to determine fitness, so the degree of bet-hedging should depend on other traits that influence whether an organism can survive until reproduction [10, 12]. For instance, the adaptive value of delaying germination depends on the risk of seed mortality of seeds stored underground and seeds freshly produced [11, 13], which can depend on seed traits, such as varying seed coat thickness. Further, the effect of variable conditions may also be mediated by traits expressed later in the life cycle, such as those that relate to the organism’s ability to tolerate stress as well as the ability to capitalize on resource bonanzas.

In desert annual plants, low-resource tolerance (i.e. stress tolerance) and resource-use capacity (i.e. the ability to capitalize on bonanzas) are likely to influence the fitness consequences of annual germination rates. One approach to assess the fitness consequences of such functional responses is through relating resource availability to two quantities: the probability of producing any offspring (reproductive success) and the number of offspring (yield upon reproductive success). The set of intercepts and slopes from these per-species relations explain performance upon low-resource availability and responsiveness to increased resource availability, respectively, and can be incorporated into models for understanding the adaptive value of germination timing. Of course, abiotic conditions, such as temperature or precipitation, are not the only sources of risk in uncertain environments. The fitness consequences of risk-spreading strategies also depend on biotic conditions, making traits that affect response to competition or other density-dependent processes important as well [14, 15]. Under density dependence, the optimal strategy depends on those employed by other individuals in the population; methods from adaptive dynamics can be used to identify evolutionarily stable strategies (ESSs) [16].

Here, we build upon a model of Gremer and Venable [11] to estimate long-term stochastic fitness in relation to germination for ten annual plants in the iconic Sonoran Desert winter annual community. We have chosen to analyze these particular species, as they are both abundant and a good representation of the variation in functional trait strategies and demography in the system [10, 11]. Here, as in Gremer and Venable [11], we use germination fractions to indicate the degree of bet hedging. Low germination fractions indicate that less seeds germinate in a given year and, instead, more remain in the seedbank. These dormant seeds serve as a “hedge” against uncertainty in any given year, so lower germination indicates higher bet hedging. Conversely, high germination fractions indicate less bet hedging, with perfect germination fractions (i.e. 100% germination) indicating that no bet hedging is occurring. Unlike Gremer and Venable [11], we directly model the effect of precipitation on plant yield for this water-limited system. In doing so, we can see how precipitation relates to the fitness, trait evolution, and bet-hedging strategies of each species. Specifically, we model the probability of reproductive success (versus reproductive failure) and per-capita yield, conditioned on reproductive success, as functions of precipitation and intraspecific density. We incorporate these relationships into a density-dependent seed bank model to predict evolutionarily stable germination strategies. To distinguish the relative importance of density, seed survival, and functional responses in driving the evolution of germination strategies, we then build a statistical regression model and use it to explore how well it predicts observed and model-predicted germination strategies.

## Model and Methods

To model the seed bank dynamics of annual plants, we adapted a seed bank model (Figure 1A) introduced by Gremer and Venable [11] and related their low-density yield to precipitation (details on the empirical data that we used to parameterise the seed bank model can be found in Appendix A3). To model low-density yield values from precipitation, we utilized a two-step Hurdle Model [17]. First, using a binomial regression, we identify the probability a germinating individual reproduces under a certain amount of precipitation (i.e. the probability that an individual crosses the “reproductive hurdle” (Figure 1B)). Then, we determine how the yield of reproducing individuals depends on this precipitation (Figure 1C) (i.e. once an individual crosses the reproductive hurdle, we determine its yield from precipitation). Incorporating our Hurdle Model into Gremer and Venable [11]’s bet-hedging, seed bank model, we calculate the evolutionarily stable germination fraction (ESS) for each of the 10 species, independently. We then use a statistical regression to estimate the relative importance of several parameters (that reflect species-specific life history traits) in governing both predicted and observed ESS germination fractions. While we use the modified bet-hedging model to predict germination values via ESS analysis, we use the statistical regression or model to assess the relative importance of different life history traits in explaining predicted and observed germination fractions.

**Figure 1:**
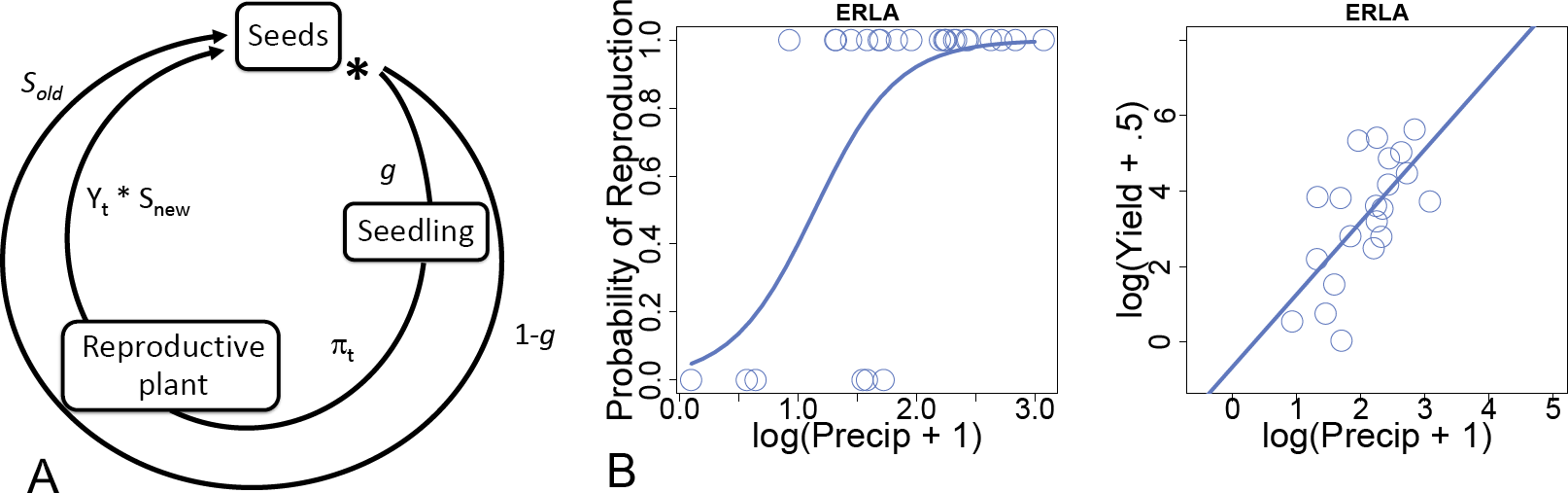
Panel A denotes the life cycle of an individual seed, as described by eqn. (1). An individual seed remains dormant with probability 1 − *g* and survives to the next Fall with probability *s*_*old*_ (outer circle). Otherwise, it germinates, becomes a reproductive plant with probability *π*_*t*_ (Panel B), and produces *Y*_*t*_ seeds per square meter (Panel C); a fraction *s*_*new*_ of *Y*_*t*_ then contributes to the following Fall (inner circle). Panel B and C represent the Hurdle Model for ERLA. Panel B shows the binomial regression for ERLA, where the open circles located on the zero and one lines represent reproductive failures and successes corresponding to precipitation values *P*_*t*_ in year *t*, and the solid, logistic curve represents the probability of reproductive success *π*_*t*_ as a function of precipitation. Panel C shows the linear regression of the post-hurdle low-density yield values *K*_*t*_ against *P*_*t*_ on a log-log scale for ERLA.

### Bet-Hedging Model

Let *n*_*t*_ denote the density of seeds in the seed bank in year *t* for a focal species. A fraction of *g* seeds germinate each year. Under low density conditions, germinating seeds contribute *K*_*t*_ seeds to the seed bank in year *t*; we refer to *K*_*t*_ as the ‘low-density yield’ in year *t*. These seeds survive to the next year with probability *s*_new_, the survival rate of fresh seeds. Negative density-dependence reduces yield by a factor 1/(1 + *agn*_*t*_), where *a >* 0 is a species-specific competition coefficient for the germinating population. A fraction of (1 − *g*) seeds remain dormant until the next year, each surviving with probability *s*_old_, the survival rate of dormant seeds (Figure 1). Under these assumptions,

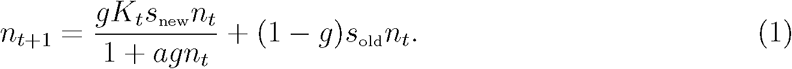

Benaїm and Schreiber [18] show that the density of seeds, given by (1), converge with probability one to a unique stationary distribution as *t* → *∞*. In particular, if the low-density per capita growth rate

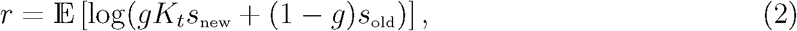

is negative, then *n*_*t*_ converges with probability one to 0. Alternatively, if this low-density per-capita growth rate is positive, *n*_*t*_ converges to a unique, positive stationary distribution 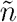. In particular, we can approximate the stationary distribution 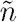 by one sufficiently long run of the model (1).

### Modeling Low-Density Yield from Precipitation: the Reproductive Hurdle

While Gremer and Venable [11] estimated *K*_*t*_ directly from the demographic data, we model how *K*_*t*_ depends on precipitation *P*_*t*_ in each year in order to analyze the effect of precipitation on seed bank dynamics. We used the same 30 years of data as Gremer and Venable [11] and obtained each year’s observed yield and precipitation values for the ten desert annuals featured in Venable [10]. Following Venable [10], a factor of 0.5 is added to yield and a factor of 1 is added to precipitation, so that the residuals of the linear regressions between log-yield and log-precipitation were approximately normally distributed. Let *N*_*t*_ = *gn*_*t*_ be the seedlings per *m*^2^ in year *t*. Then the observed yield is 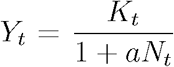, where *a* and *K*_*t*_ are the measured competitive coefficient and measured low-density yield for year *t*, both calculated by Gremer and Venable [11].

For our two-step Hurdle Model relating low-density yield to precipitation, we model the probability of reproductive failure (*K*_*t*_ *<* 1) or success (*K*_*t*_ *≥* 1) with a binomial regression (Figure 1B). Specifically, we keep track of each species’ reproductive failures and successes and their corresponding log-precipitation values, all of which were measured within the 30 years of yield and precipitation data. We then binomially regress these failures and successes against log-precipitation, the regression being the reproductive hurdle or probability an individual reproduces as a function of log-precipitation. For the subset of years that a species experienced reproductive success (i.e. cleared the hurdle), we linearly regress non-zero, low-density yield *K*_*t*_ against precipitation *P*_*t*_ on the log scale (Figure 1C): log(*K*_*t*_) = log(*α*_2_) + *β*_2_ log(*P*_*t*_), where *P*_*t*_ is the measured precipitation for year *t*. The reproductive hurdle and the post-hurdle regressions define a two-step process relating the low-density yield *K*_*t*_ to the precipitation *P*_*t*_:

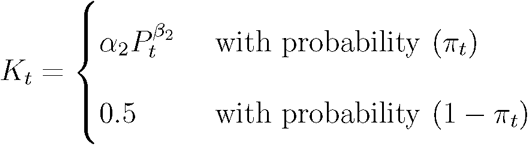

where the probability of crossing the reproductive hurdle is:

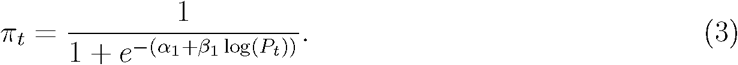

We call *α*_1_ the reproductive intercept, *β*_1_ the reproductive slope, *α*_2_ the log-yield intercept, and *β*_2_ the log-yield slope. Larger reproductive intercepts and reproductive slopes correspond to higher probabilities of reproductive success in low and high precipitation years, respectively. Larger log-yield intercepts and log-yield slopes correspond to higher yield in low and high precipitation years, respectively. Larger log-yield intercepts represent increased ability to reproduce under low resource availability (i.e. tolerance to low water availability), whereas larger values for log-yield slopes represent increased responsiveness to resources and ability to capitalize on wet years for reproduction.

### Predicting ESS Values

To identify the ESS for the germination fraction *g*, consider a “mutant” population at very low density ñ with a different germination strategy, 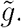. At low densities, this mutant population has a negligible feedback on the resident population and itself, but is influenced by the resident. Hence, its population dynamics are initially approximated by

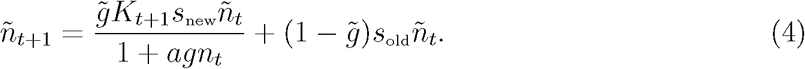

The success or failure of the invasion of these mutants is determined by their stochastic growth rate,

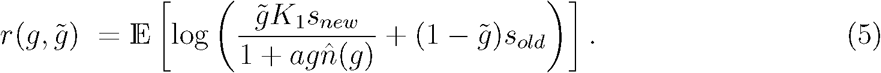

If 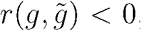, the invasion fails and if 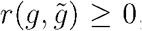, it succeeds [19]. A germination strategy *g* is an ESS if 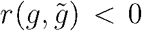 0 for all 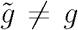 (i.e. no other strategy can invade). A necessary condition for an ESS with 0 *< g <* 1 is that 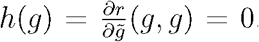. Since 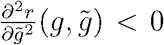 for all 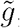, if such a *g* is found, it is unique. Thus to find the ESS, we solve for the root of *h*(*g*) strictly between 0 and 1 (see Appendix A1).

## Results

### Reproductive Hurdle Model

Species were significantly different in their reproductive and log-yield intercepts for both the binomial regression modeling reproductive success, and the post-hurdle model. For a common slope, binomial sub-model with solely log-precipitation and species as predictors, a pairwise Wald’s test detected differences between multiple pairs of reproductive intercepts. In particular, EVMU significantly differed from ERCI, PERE, PLIN, PLPA, and SCBA (*p* < .05); STMI significantly differed from PERE and SCBA (*p <* .05) (see Table 1 for species’ full nomenclature). The rest of the pairs were not significantly different, according to the pairwise Wald’s test. A post-hoc Tukey test with a Bonferroni correction also did not reveal significant differences for any pairing. We compared the common slope sub-model to a common intercept sub-model (the interaction and log-precipitation as predictors). The common slope, submodel was the winning model in AIC, BIC, and log-likelihood comparisons and was not significantly different from the full binomial model. The binomial fit (the coefficients coming from the full binomial model) for *Eriophyllum lanosum* (ERLA) is shown in Figure 1B.

**Table 1:** This table contains (from left to right) values for seed survivorship in the seed bank, seed survivorship of fresh seeds, reproductive intercepts, reproductive slopes, log-yield intercepts, log-yield slopes, and competition factors. Superscripted dots and stars next to *α* and *β* values indicate how significantly different these parameters are from 0: .1^•^, .05^∗^,.01^∗∗^, and .001^∗∗∗^.

| Species | $s_{old}$ | $s_{new}$ | $\alpha_1$ | $\beta_1$ | $\alpha_2$ | $\beta_2$ | $a$ |
| --- | --- | --- | --- | --- | --- | --- | --- |
| *Monoptilon belliodes* (MOBE) | .273 | .102 | -3.31<sup>•</sup> | 2.74<sup>*</sup> | 1.73 | 0.79 | .0163 |
| *Erodium cicutarium* (ERCI) | .429 | .132 | -1.55 | 1.97<sup>*</sup> | 0.02 | 1.25<sup>**</sup> | .0351 |
| *Stylocline micropoides* (STMI) | .458 | .145 | -1.78 | 1.46<sup>*</sup> | -1.67 | 2.33<sup>***</sup> | .0156 |
| *Schismus barbatus* (SCBA) | .512 | .150 | -2.03 | 3.10<sup>*</sup> | -0.54 | 2.38<sup>***</sup> | .0112 |
| *Eriophyllum lanosum* (ERLA) | .532 | .153 | -3.15<sup>•</sup> | 2.86<sup>*</sup> | -0.73 | 1.95<sup>***</sup> | .0114 |
| *Plantago patagonica* (PLPA) | .550 | .160 | -2.18 | 2.43<sup>*</sup> | 1.45 | 0.84 | .0069 |
| *Erodium texanum* (ERTE) | .577 | .170 | -1.29 | 1.43<sup>•</sup> | -0.07 | 1.36<sup>*</sup> | .0532 |
| *Plantago insularis* (PLIN) | .596 | .168 | -1.42 | 2.14<sup>*</sup> | 1.52 | 0.89<sup>•</sup> | .0380 |
| *Pectocarya recurvata* (PERE) | .627 | .173 | -0.10 | 1.43 | 0.93 | 1.44<sup>***</sup> | .0167 |
| *Evax multicaulis* (EVMU) | .828 | .214 | -3.86<sup>*</sup> | 2.36<sup>*</sup> | 1.58 | 1.30<sup>*</sup> | .0119 |

A common intercept, post-hurdle model showed SCBA’s log-yield slope significantly differed from ERCI, ERLA, ERTE, MOBE, PLIN, PLPA, and STMI; PERE significantly differed from ERCI and ERTE; EVMU significantly differed from ERCI, ERTE, PLPA, and STMI (*p <* .05). Non-listed pairs of log-yield slopes were not significantly different from one another. In addition, a post-hoc Tukey test with a Bonferroni correction revealed significant differences between EVMU and ERCI, ERTE and SCBA, and ERCI and SCBA (*p <* .05). No other pairs were significantly different. In contrast, a common slope submodel also yielded significant intercepts but was not selected after comparing AIC, BIC and log-likelihood values to the common intercept sub-model. The common intercept sub-model was also found to not be significantly different from the full, linear model. The post-hurdle fit (the coefficients coming from the full, linear model) for ERLA is shown in Figure 1C.

### Predicted ESS Values

With the purpose of estimating ESS values as accurately as possible, we used the full models’ intercepts and slopes, unique for each species, for both reproductive success and post-hurdle yields. We have incorporated the resulting reproductive and log-yield intercepts and slopes, survival rates within the seed bank, and the competitive responses into one table (Table 1). Predicted ESS values from the full models corresponded well with observed germination fractions; our ESS values explain 69% of the variance (adjusted *R*^2^) in observed germination fractions (on the logit-logit scale), although our ESS values typically overestimate these germination fractions (Figure 2).

**Figure 2:**
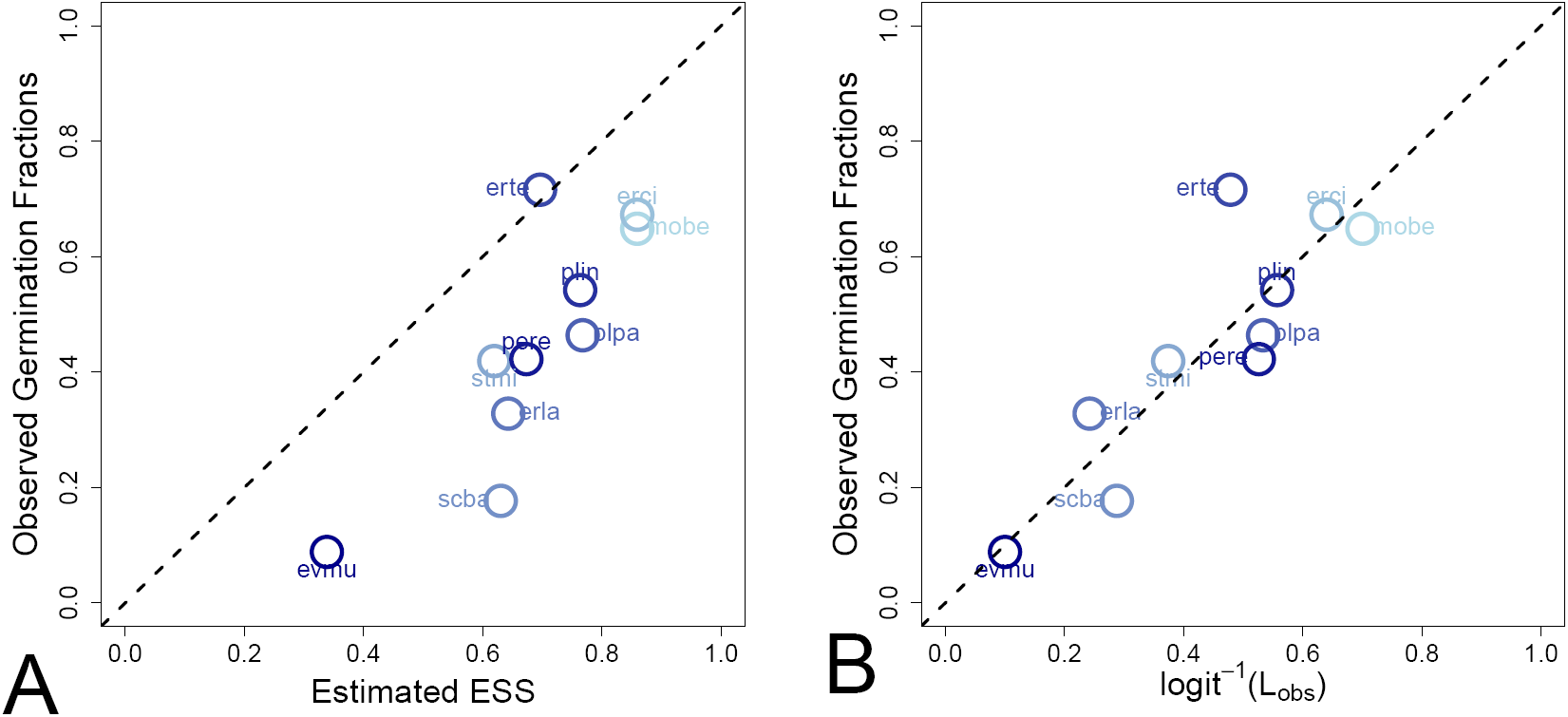
Panel A: the observed germination fractions against model-estimated ESS values for each species. Panel B: germination fractions against germination estimates found by taking the inverse-logit of *L*_*obs*_. Points lying on the dotted line represents equality between predicted ESS values and observed germination fractions. Points have been colored from light blue (low seed survival) to dark blue (high seed survival).

### Factors Shaping Bet Hedging

To explore the effect of each trait on ESS values and observed germination rates, we also examined how well our model-estimated ESS values and observed germination fractions could be estimated by parameters of Table 1 on the logit scale. We used our ANOVA – along with the non-dimensionalization of equation (1) (see Appendix A2) – to reduce the potential predictors to reproductive intercepts, log-yield slopes, and survival rates of dormant seeds. The resulting linear regression

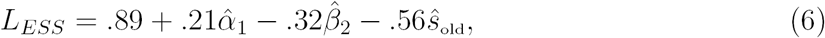

explains 90% (adjusted *R*^2^) of the variance in the predicted ESS values, where the hatted variables denote the standardized predictor variables. Survival rates of dormant seeds have the most impact on variation in ESS values, followed by log-yield slopes and reproductive intercepts. Finally, we analyzed the amount of variance in observed germination fractions explained by these three parameters. The resulting linear regression

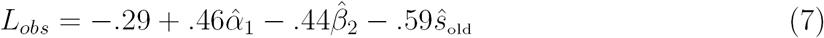

explains 69% (adjusted *R*^2^) of the variance in the observed germination fractions (Figure 2). Seed survival rates again have the largest negative effect on the germination rates. In contrast, reproductive intercepts have a much larger positive effect on observed germination rates than they did in the model-predicted ESS values. Log-yield slopes had a much stronger negative impact on observed germination rates than the model-predicted ESS values. Both regressions show that germination fractions are lower for species that have larger survival rates of seeds in the seed bank (*s*_old_), higher for species with greater reproductive success in dry years (*α*_1_), and lower for species that capitalize on wet years upon reproductive success (*β*_2_).

## Discussion

Our approach, which explicitly incorporates an aspect of environmental variation, and species-specific responses to that variation, into models for understanding the factors that shape optimal strategies, illuminates life-history evolution in variable environments and is likely to be useful in many other systems. The ESS model predicts germination fractions based on uncertainty in precipitation, whereas the statistical model indicates the relative importance of survival rates, reproductive success, and precipitation-dependent yield. Most important is seed survivorship: the safer it is to be in the seed bank, the more beneficial it is to bet-hedge, a result consistent with classical bet-hedging theory [9, 13]. Equally as important as the precipitation-dependent yield (but less so than seed survivability) is the sensitivity of reproductive success in low rainfall years. Higher reproductive success in dry years selects for desert annuals to bet-hedge less. However, higher rates of seed production in wet years push annuals to bet-hedge more, which could be attributed to higher rates of seed production in wet years being linked to lower rates of seed production in dry years.

Indeed, empirical observations in this system indicate that some species are better able to capitalize on favorable conditions than others. For instance, Angert et al. [20] experimentally demonstrated differences in leaf area and biomass allocation in response to precipitation pulses between two species of winter annuals. The more stress tolerant species (*Pectocarya recurvata*) was less able to capitalize on the resource pulse than the other, less tolerant, species (*Stylocline micropoides*). Similarly, Kimball et al. [21] showed that species with high relative growth rates (RGR) and low water-use efficiency (WUE) were favored in years with large, and more frequent rain events, using 25 years of climate and fitness data for this winter annual system. These functional tradeoffs also relate to germination biology; resource acquisitive species with low germination fractions also have germination physiologies cued to slower germination and in a narrower range of conditions [22]. In other words, they have physiologies tuned to slower and more cautious germination, but also have less tolerant, resource acquisitive traits once established. Thus, functional responses included in our ESS analyses strongly affect long-term variation in fitness and germination in response to precipitation in this system.

Our ESS estimates explained approximately 29% more variance in germination fractions than a linear regression on Gremer & Venable’s predicted values; although, we consistently overestimate the observed germination fractions, whereas their predictions are a mixture of under- and over-predicted values with smaller summed squared-residuals (Figure 2A) [11]. Our over-estimates might stem from all the possible risks or factors that may select for greater bet-hedging, e.g. seed predation, interspecific competition, variable temperature, and disease [23–29], which were not included in our model. Furthermore, 30 years of precipitation data might not fully capture the long-term variation in precipitation that has shaped the evolution of germination timing in these species.

Our analyses also indicate differences in what is driving predicted ESS values versus observed germination rates. While the general intercept, reproductive yield, and seed survival rates within the seed bank affect predicted ESS values (see equation 6), reproductive success is a better predictor of observed germination fractions (see equation 7). The general intercepts of the two regressions account for the largest difference in fit between the statistical model and the ESS models’ predicted germination fractions. This major difference partially stems from the variance in our predicted yield values generally having lower variance than that of the estimated low density yield values of Gremer & Venable [11]. Larger variation in our yield values would select for more bet-hedging (lower ESS estimates) for all ten species, and consequently, subtract from the general intercept to equation 6. The second largest difference is that of the coefficient of the *α*_1_ term, or standardized reproductive intercept. Since reproductive success is also determined solely from precipitation in our model, the difference in coefficient magnitudes may be highlighting our inability to capture variation in reproductive success from rainfall alone. Thus, the statistical model suggests that the variance in both yield and reproductive success is not completely explained by the variation in the 30 years of precipitation. Other factors such as temperature may also play a role in mediating responses to water availability, particularly in this system, and could be incorporated in future models [21, 30, 31].

As a model for displaying the relative importance of survival rates, reproduction rates, and conditional seed production rates, the statistical regression for observed germination fractions suggests that we are not fully capturing variance in yield and reproductive values. Thus, discerning these biological uncertainties and modeling them will be an important step toward understanding the reasons why desert annuals hedge their bets as much as they do. Moreover, our ESS model can be a significant springboard to predicting germination rates for desert annuals: not only can one fine-tune the ESS estimates we have predicted by incorporating variable factors such as temperature and predation, one may also use this same ESS machinery to predict germination rates of desert annuals outside of the ten studied here. Finally, as 30 years of data did not capture the whole picture in mean and variation in precipitation, it will be equally important to investigate how differences in either of these affect evolutionarily stable strategies.

## Data Accessibility

All measured quantities for this study can be found at Dr. Lawrence Venable’s Desert Annual Archive (http://www.eebweb.arizona.edu/faculty/venable/LTREB/LTREB%20data.htm).

## Authors’ Contributions

W.S.C. and S.J.S. conceived of the study and developed the model and analytical methods. All authors gave feedback on the model and methods and were involved in the interpretation of the results. W.S.C. carried out the statistical analysis and model simulations. W.S.C. and J.R.G. wrote the manuscript. All authors contributed to content and editing of the manuscript.

## Competing Interests

The authors have no competing interests.

## Funding

This work was supported by the NSF (IOS 1456724 to A. Sih and DMS 1313418 & 1716803 to S.J. Schreiber).

## Acknowledgments

We thank Dr. Larry Venable for the demography and precipitation data.

## References

[1] M. Slatkin. Competition and regional coexistence. Ecology, 55(1):128–134, 1974.

[2] J. Seger and H.J. Brockmann. What is bet-hedging? Oxford Surveys in Evolutionary Biology, 4:182–211, 1987.

[3] S.J. Schreiber. Unifying within- and between-generation bet-hedging theories: an ode to J.H. Gillespie. The American Naturalist, 186(6):792–796, 2015.

[4] S. Tuljapurkar. Delayed reproduction and fitness in variable environments. Proceedings of the National Academy of Sciences, 87(3):1139–1143, 1990.

[5] N.G. Hairston Jr and W.R. Munns Jr. The timing of copepod diapause as an evolutionarily stable strategy. The American Naturalist, 123(6):733–751, 1984.

[6] F. Menu. Strategies of emergence in the chestnut weevil curculio elephas (coleoptera: Curculionidae). Oecologia, 96(3):383–390, 1993.

[7] M.L. Crump. Variation in propagule size as a function of environmental uncertainty for tree frogs. The American Naturalist, 117(5):724–737, 1981.

[8] A. Charpentier, M. Anand, and C.T. Bauch. Variable offspring size as an adaptation to environmental heterogeneity in a clonal plant species: integrating experimental and modelling approaches. Journal of Ecology, 100(1):184–195, 2012.

[9] D. Cohen. Optimizing reproduction in a randomly varying environment. Journal of Theoretical Biology, 12(1):119–129, 1966.

[10] D.L. Venable. Bet hedging in a guild of desert annuals. Ecology, 88(5):1086–1090, 2007.

[11] J.R. Gremer and D.L. Venable. Bet hedging in desert winter annual plants: optimal germination strategies in a variable environment. Ecology Letters, 17(3):380–387, 2014.

[12] J. M. McNamara, Z. Barta, M. Klaassen, and S. Bauer. Cues and the optimal timing of activities under environmental changes. Ecology Letters, 14(12):1183–1190, 2011.

[13] S. Ellner. ESS germination strategies in randomly varying environments. I. Logistic-type models. Theoretical Population Biology, 28(1):50–79, 1985.

[14] S. Ellner. Alternate plant life history strategies and coexistence in randomly varying environments. In Theory and Models in Vegetation Science, pages 199–208. Springer, 1987.

[15] M.J. Groom. Allee effects limit population viability of an annual plant. The American Naturalist, 151(6):487–496, 1998.

[16] O. Diekmann. A beginners guide to adaptive dynamics. Summer School on Mathematical Biology, pages 63–100, 2002.

[17] J. Mullahy. Specification and testing of some modified count data models. Journal of Econometrics, 33(3):341–365, 1986.

[18] M. Benaïm and S.J. Schreiber. Persistence of structured populations in random environments. Theoretical Population Biology, 76(1):19–34, 2009.

[19] P. Chesson and S. Ellner. Invasibility and stochastic boundedness in monotonic competition models. Journal of Mathematical Biology, 27:117–138, 1989.

[20] A.L. Angert, J.L. Horst, T.E. Huxman, and D.L. Venable. Phenotypic plasticity and precipitation response in sonoran desert winter annuals. American Journal of Botany, 97(3):405–411, 2010.

[21] S. Kimball, J.R. Gremer, A.L. Angert, T.E. Huxman, and D.L. Venable. Fitness and physiology in a variable environment. Oecologia, 169(2):319–329, 2012.

[22] Z. Huang, S. Liu, K.J. Bradford, T.E. Huxman, and D.L. Venable. The contribution of germination functional traits to population dynamics of a desert plant community. Ecology, 97(1):250–261, 2016.

[23] M. Kistenmacher and J.P. Gibson. Bet-hedging against larval herbivory and seed bank mortality in the evolution of heterocarpy. American Journal of Botany, 103(8):1383–1395, 2016.

[24] J.L. Horst and D.L. Venable. Frequency-dependent seed predation by rodents on Sonoran Desert winter annual plants. Ecology, 2017.

[25] S. Kalisz. Variable selection on the timing of germination in *Collinsia verna (Scrophulariaceae)*. Evolution, 40(3):479–491, 1986.

[26] K. Tielbörger and A. Valleriani. Can seeds predict their future? Germination strategies of density-regulated desert annuals. Oikos, 111(2):235–244, 2005.

[27] A.R. Dyer, A. Fenech, and K.J. Rice. Accelerated seedling emergence in interspecific competitive neighbourhoods. Ecology Letters, 3(6):523–529, 2000.

[28] M.K.J. Ooi, T.D. Auld, and A.J. Denham. Climate change and bet-hedging: interactions between increased soil temperatures and seed bank persistence. Global Change Biology, 15(10):2375–2386, 2009.

[29] J.W. Dalling, A.S. Davis, B.J. Schutte, and A.A. Elizabeth. Seed survival in soil: interacting effects of predation, dormancy and the soil microbial community. Journal of Ecology, 99(1):89–95, 2011.

[30] J.R. Gremer, S. Kimball, A.L. Angert, D.L. Venable, and T.E. Huxman. Variation in photosynthetic response to temperature in a guild of winter annual plants. Ecology, 93 (12):2693–2704, 2012.

[31] T.E. Huxman, S. Kimball, A.L. Angert, J.R. Gremer, G.A. Barron-Gafford, and D.L. Venable. Understanding past, contemporary, and future dynamics of plants, populations, and communities using Sonoran Desert winter annuals. American Journal of Botany, 100(7):1369–1380, 2013.

